# Both social environment and chronological age shape the physiology of ant workers

**DOI:** 10.1101/2022.06.13.495886

**Authors:** Martin Quque, Charlotte Brun, Claire Villette, Cédric Sueur, François Criscuolo, Dimitri Heintz, Fabrice Bertile

## Abstract

Position within the social group has consequences on individual lifespans in diverse taxa. This is especially obvious in eusocial insects, where workers differ in both the tasks they perform and their aging rates. However, in eusocial wasps, bees and ants, the performed task usually depends strongly on age. As such, untangling the effects of social role and age on worker physiology is a key step towards understanding the coevolution of sociality and aging. We performed an experimental protocol that allowed a separate analysis of these two factors using four groups of black garden ant (*Lasius niger*) workers: young foragers, old foragers, young nest workers, and old nest workers. We highlighted age-related differences in the proteome and metabolome of workers that were primarily related to worker subcaste and only secondarily to age. The relative abundance of proteins and metabolites suggests an improved xenobiotic detoxification, and a fuel metabolism based more on lipid use than carbohydrate use in young ants, regardless of their social role. Regardless of age, proteins related to the digestive function were more abundant in nest workers than in foragers. Old foragers were mostly characterized by weak abundances of molecules with an antibiotic activity or involved in chemical communication. Finally, our results suggest that even in tiny species, extended lifespan may require to mitigate cancer risks. This is consistent with results found in eusocial rodents and thus opens up the discussion of shared mechanisms among distant taxa and the influence of sociality on life history traits such as longevity.

## 1. Introduction

The place that an individual holds in the social network of the group has emerged as a strong explanatory factor of aging in various taxa (reviewed in 1): birds (2, 3), insects (4–6), and mammals (7) including humans (7–9). To better understand such a well conserved evolutionary relationship between sociality and aging, the use of additional key species that evolved remarkable social organizations will make it possible to test specific hypotheses experimentally and conduct studies faster than in humans. Some bee, ant, wasp, and termite species are known for their well-established social structure in which each individual performs tasks specific to the behavioral caste to which it belongs. Such a social division of labor has led to the naming of these insects as eusocial insects. In recent years, many studies have shown that eusocial insects are relevant models to understand the mechanistic basis of lifespan diversity (10, 11). For example, reproductive individuals in social insects may live up to 100 times longer than individuals of solitary insect species (12). However, the mechanisms usually known to be associated with aging in mammals or birds sometimes show either no or opposite relationships in eusocial insects. In fact, neither antioxidant enzymes (13–15), nor oxidative damage (reviewed in 16) appeared so far to be strongly associated with lifespan in social insects. Moreover, telomere length or their attrition rate are identified as reliable indicators of life expectancy in mammals and birds *(e.g.* 17–19). However, in the black garden ant *(Lasius niger),* telomeres are of similar length in workers and queens, while the latter live ten times longer (20).

Interpreting the results of such studies is sometimes complicated by the fact that the social role of a worker subcaste and its age are intimately related. The youngest workers most often perform tasks inside the nest (*e.g.*, feeding queen, building the nest) and the oldest ones perform tasks outside the nest (*e.g.*, food provisioning, waste management). Through experimental demographic manipulations, old workers can be forced to resume nest worker tasks and young workers to forage, opposing the classical pattern. By using this approach, a few studies have tried to disentangle the respective influences of chronological age and behavior on the physiology of eusocial insect workers (21–24). Some of those studies found that the social role was better than chronological age to explain the differences in gene expression or proteome (21, 22, 24). Conversely, despite significant differences in behavior, morphology, and longevity between the most extreme social castes (i.e. queen and workers), transcriptomics comparison in *Lasius niger* showed that the divergence in gene expression was better explained by age than behavior (25). Finally, some metabolic characteristics turn to be found varying in both an age-and behavior-dependent manner, like the concentration of brood pheromones involved in caste maturation (26). Thus, there is so far no consensus on the respective importance of age and social role effects on the physiology of social insects.

To address this question, we successively sampled workers from colonies of black garden ants. The first sampling occurred at the time of colony foundation where both worker subcastes had young individuals. One year later, we sampled the same colonies a second time. As the eggs were regularly removed before hatching, ants of both worker subcastes were one year older. Thanks to this unprecedented protocol, we have obtained old nest workers that were very unlikely to have ever been foragers and thus retained no potential molecular trace of past foraging activities. We then jointly analyzed the metabolome and proteome to get a comprehensive picture of physiological changes. By separating the effects of social role and age on physiology, this study aims to better understand the mechanisms of aging and to what extent the social environment can influence it.

## 2. Results and Discussion

The combined mass spectrometry analyses (LC-MS/MS) detected 712 metabolites and 1719 proteins (**Figure S1**). Original data sets are available in electronic supplementary material (**Table S1-S3**). The tables used for the fold-change (FC) analysis, cleaned from metabolites or proteins found in less than 3/5 samples in a group, are also available online (**Tables S4 and S5)**. Supplementary materials also encompass the statistics summary (FC, FDR, class, biological processes) of these analyses (**Tables S7 and S8**), as well as the references used to link analytes (proteins + metabolites) to the biological processes mentioned below. We found 16 proteins and 11 metabolites completely absent from at least one worker subcaste (**Table S9 and S10).** No functional annotation and no relevant studies were found to help us to attribute a clear biological meaning to them. Consequently, absent proteins and metabolites will not be further discussed thereafter.

We drew heat maps from the metabolomics and proteomics data to picture how the four experimental groups (Y.F: young foragers, O.F: old foragers, Y.NW: young nest workers, O. NW: old nest workers) clustered together. The metabolomics-based clustering (**Figure 1**) shows that the four experimental groups have a weaker intra-group variability than inter-group variability and the O.F group differs the most when compared to the three other groups. Still, the two groups of nest workers (Y.NW and O.NW) are closer to each other than to Y.F, indicating that the behavior has a stronger impact than age on individual metabolome. On the contrary, the proteomics-based clustering (**Figure 2**) assorts samples according to the age (old or young) but independently of the worker subcaste (NW or F). As the relative importance of worker subcaste and age may depend on the biological processes studied, it is understandable that past studies found conflicting results about the role of age and social role (24, 25).

**Figure 1.**
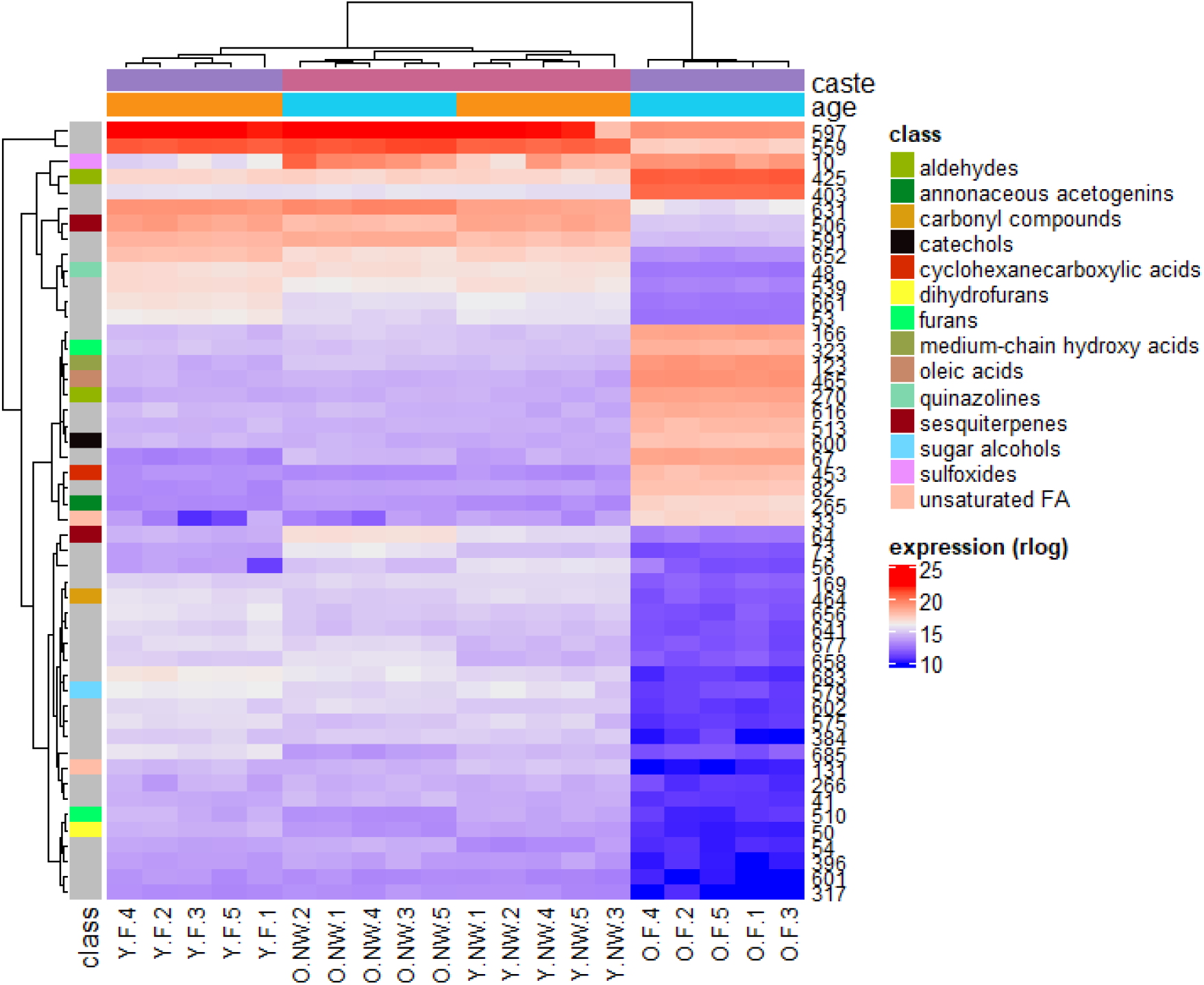
Heat map of the 50 most expressed metabolites amongst our four experimental groups in *L. niger.* At the top, colours indicate the age (**orange**: young, **light blue**: old) and caste (**purple**: foragers **pink**: nest-workers,). The left column indicates the metabolite class. Grey means a NA value. At the bottom are the sample identifiers (F: forager, NW: nest-worker, Y: young, O: old). For a given metabolite, the right column indicates its ID rather than the full name for legibility reasons (correspondence is indicated in every table provided). All metabolites presented here have an FDR < 0.05 and |fold-change| > 2.

**Figure 2.**
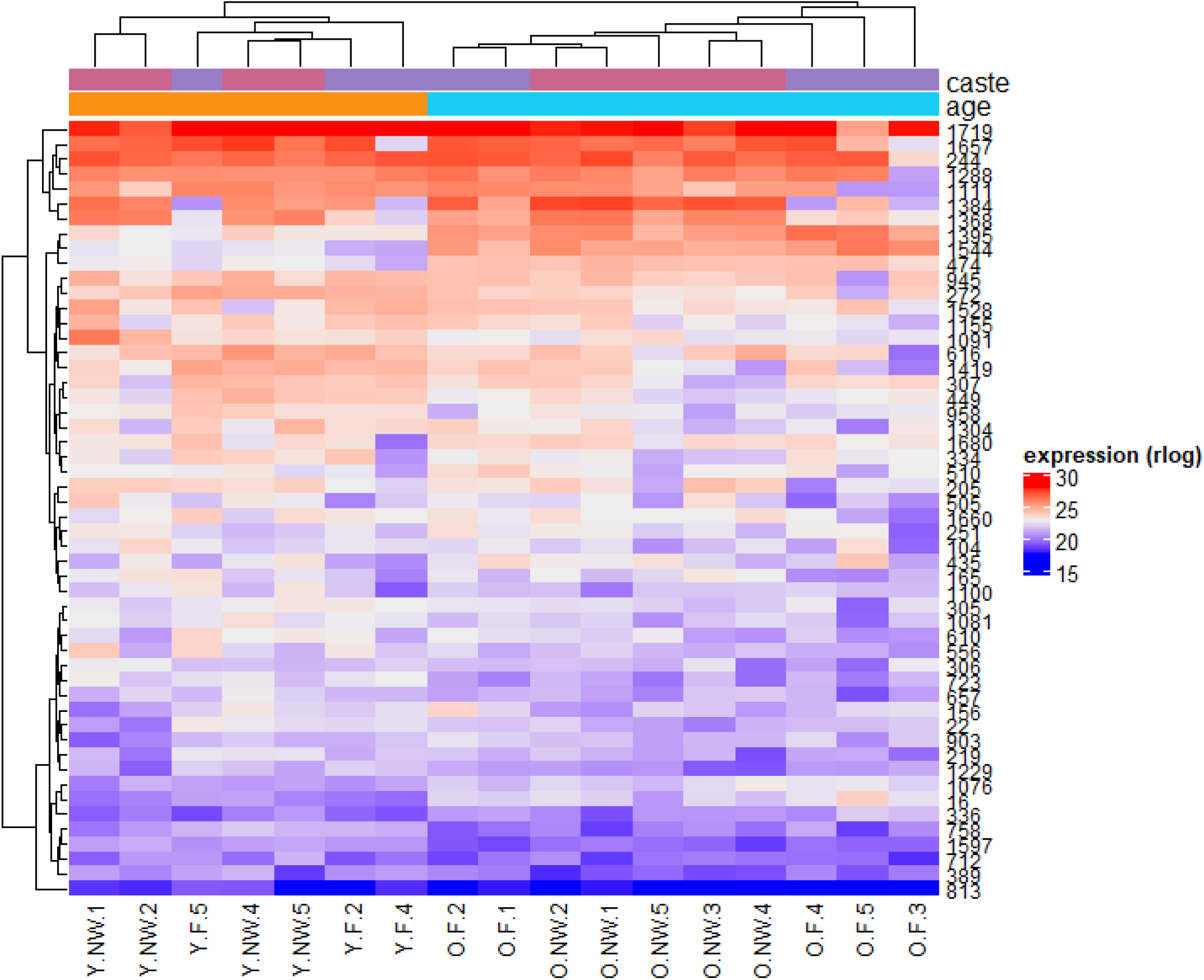
Heat map of the 50 most expressed proteins amongst our four experimental groups in *L. niger.* At the top, colours indicate the age (**orange**: young, **light blue**: old) and caste (**pink**: nest-workers, **purple**: foragers). At the bottom are the sample identifiers (F: forager, NW: nest-worker, Y: young, O: old). For a given protein, the right column indicates its ID rather than the full name for legibility reasons (correspondence indicated in every table provided). All proteins have an FDR< 0.05 and |fold-change| > 2.

### 2.1. Old ants show a poorer somatic maintenance

Compared to O.NW, Y.NW had greater quantities of cytochrome P450 4g15, belonging to the CYP4 subfamily of cytochrome P450. Although potentially to a lesser extent than CYP1, 2 and 3 proteins (27), CYP4 have been found to be linked to the degradation of toxic xenobiotics (28–30). Similarly, Y.F compared to O.F had greater quantities of this same protein, as well as transferrin, which limits bacterial infection through iron-chelation (31). The metabolome analysis was consistent with the proteome analysis as it found greater quantities in Y.F than in O.F of metabolites associated with xenobiotic degradation, *e.g.*, 1-methoxyphenanthrene, cyclohexanone (KEGG maps 00930 and 00624); and immunity, *e.g.*, cyclohexane undecanoic acid, armillarin (32, 33). Besides, O.F had larger amounts of four metabolites associated with oxidative damage *(e.g,* 3,4-dimethoxybenzoic acid, robustocin), and only one of such metabolites (psilostachyin) had a larger amount in Y.F. Transferrin, which was found in greater quantities in Y.F (vs O.F) and Y.NW (vs O.NW), also has a protective role against oxidative stress (34, 35). Taken together these data depict the young ant workers as more likely to protect their organism integrity from both external (pathogens, pollutants) and internal (oxidative stress) threats. When compared to O.F, O.NW exhibit larger quantities of metabolites related to immunity (*e.g.*, thiophene, nebularine) and xenobiotic detoxification (cyclohexanone). Poorer somatic maintenance might therefore be a marker of senescence in worker ants, and old foragers might be the subcaste that suffers the most from it. This result fuels the adversarial debate between studies showing a cumulative deterioration of the organism with age in social insects and others showing the opposite, making it difficult in the current state of knowledge to describe a consistent model (16). Furthermore, these results show that both age and social role participate in the determination of the individual phenotype.

Finally, these results have to be viewed in relation to those presented below on cancer-related molecules. Indeed, it is known that immunity has anti-cancer roles (36), and several of our metabolites and proteins are known to be associated with immunity-related defense against cancer (*e.g.*, methylfurfuryl alcohol, armillarin, psilostachyin, thiophene). This exemplifies the interconnection between anti/pro-aging processes in a large variety of organisms.

### 2.2. Ants also experience a metabolic shift when aging

The organism’s source of energy (lipids or carbohydrates) has been shown to affect longevity. For example, a previous study in black garden ants has positively associated a high body fat content with a greater probability of survival of workers under stress (23). In honey bees, the greater longevity of queens has been found positively associated with a low polyunsaturated lipids/monounsaturated lipids ratio (37). A similar relationship has been reported in long-lived birds and mammals (38, 39). All these observations suggest that lipid metabolism is crucial in determining either survival or lifespan. However, our study suggests a metabolic shift with age, from lipid to carbohydrate use in ant workers. Indeed, compared to O.F, Y.F had greater amounts of two metabolites related to glycerophospholipid metabolism (lysoPE(0:0/18:1(11Z)) and lysolecithin), and eight related to unsaturated fatty acid metabolism (e.g., (1RS,2RS)- guaiacylglycerol 1-glucoside, 14,16-nonacosanedione, 6,8-tricosanedione). Other metabolites related to lipid assimilation and metabolism in general, not only unsaturated fatty acids, were found more abundant in Y.F *(e.g.,* ximenic acid, (E)-2-tetracosenoic acid, 9-Decenoylcarnitine).

Besides, we found greater abundances in Y.F vs O.F of proteins involved in lipid metabolism (e.g., acetyl-cytosolic). On the other hand, O.F (*vs* Y.F) showed greater abundances of many proteins involved in carbohydrate metabolism (*e.g.*, glucosidase, maltase 1, trehalose transporter tret1-like) but no protein related to lipid metabolism were found more abundant in this group. Similarly, we found a higher abundance for proteins involved in carbohydrate metabolism (*e.g.*, alpha-like protein) in O.NW vs Y.NW. Interestingly, we found greater abundances of metabolites in connection with lipid metabolism ((9Z,11E)-(13S)-13-hydroperoxyoctadeca-9,11-dienoate, and a C16 ceramide (d18:1/16:0) in O.NW compared to O.F. Similarly, larger amounts of proteins involved in lipid storage (lipid storage droplets surface-binding protein 2) were observed in O.NW *vs.* O.F. Hence, the apparent metabolic shift from lipid to carbohydrate metabolism as workers age appeared more pronounced in foragers, which is the worker subcaste that ages the fastest (40, 41). In humans too, when we age, lipids are less used as an energy source (42). Less efficient use of lipids might therefore be a signature of aging common to phylogenetically very far species. Still, we cannot rule out the possibility that this may also be a side-effect accompanying other functional changes with age. For instance, the relative abundance of few molecules suggested a less active immunity or chemical communication in O.F. than Y.F (see above), and these functions are lipid consuming (43–46), which could explain a slowing down of lipid use in O.F. Whether the observed metabolic shift has a causal effect on workers’ aging rate, for instance *via* deleterious impact due to carbohydrate use as fuel (47–49), needs further experimental studies.

### 2.3. The consequences of social division of labor on digestion and communication

When comparing the proteome of nest workers with that of foragers (Y.NW *vs.* Y.F and O.NW *vs.* O.F), we systematically found the arylphorin subunit alpha in greater amount for nest workers, independently of age. This protein stimulates stem cells in the midgut and notably allows its regeneration after stress (50–52). The importance of digestive function in *Lasius niger* nest workers seems strong since we already found it in two previous studies (53, 54). To our knowledge, no other study experimentally addressed this question in adult nest workers. Two non-mutually exclusive hypotheses can be formulated to explain this phenotype: (i) the excess of food is stored and pre-digested by nest workers to make it quickly available for the rest of the colony in case of a food shortage; (ii) this pre-digested food would allow queens to process food more efficiently, by lowering energy investment in their own digestive system. The mechanism proposed in the latter hypothesis might form an energy trade-off solution explaining the unexpected equation of high reproduction and high longevity in ant queens.

When comparing O.F to Y.F (age effect) or O.NW (social role effect), among eight compounds involved in chemical communication *(e.g.,* tetracosanedioic acid, oleamide, hexadecanedioic acid, 2-[(methylthio)methyl]-2-butenal), only one (octanal) was found more abundant in O.F. This suggests that regardless of age and social role, O.F appeared to communicate much less with their congeners than do the ants in the other groups. Besides, the over-representation of oleic acid and derivatives in O.F (**Figure 1 and 3**) could be interpreted as a possible mechanism for members of the colony to detect senescent individuals. Indeed, oleic acid, linoleic acid and their derivatives are known to be secreted in a very conservative manner within insect taxa, including ants, when they die (55–57). It is known that this signal helps ants to recognize a corpse and is involved in a broad diversity of corpse management behaviors (58, 59), such as removal, cannibalism, burial (57, 58, 60, 61). It is also known that ants practice social immunity, for instance, by reducing interactions with an individual recognized as sick (62, 63). Hence, oleic acid-derived signals may allow behavioral adaptions in social interactions, *e.g.,* avoiding interactions with O.F, which potentially care more germs and might have fewer antibiotic metabolites to tackle them according to our previously discussed results. It has also been shown that sick ants isolate themselves before dying (64, 65). Though causality is difficult to assess here, both cases (ignored by others or self-isolation) would lead to the social isolation of O.F and are consistent with a lower synthesis of communication molecules.

**Figure 3.**
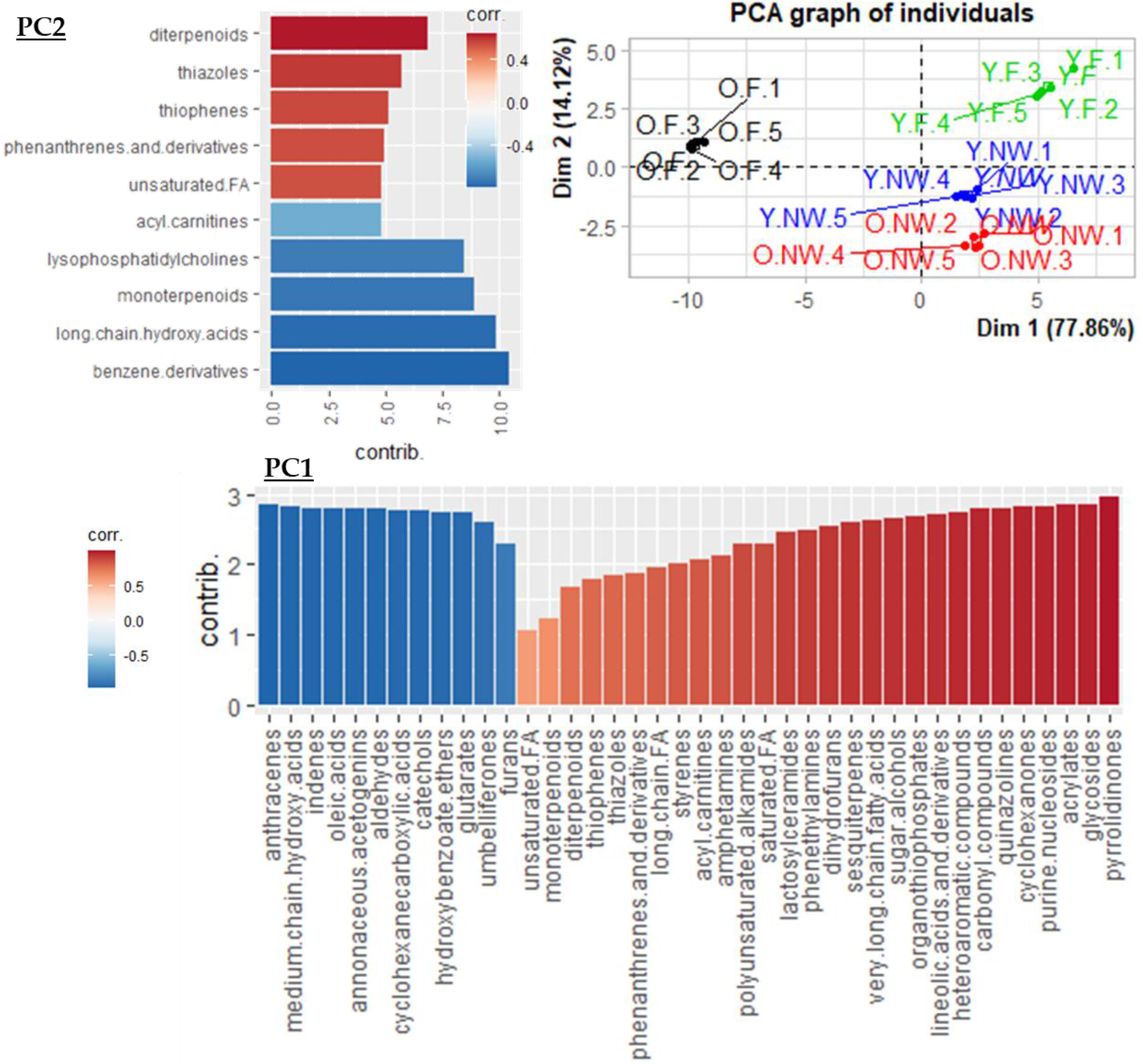
Classification-based PCA (PC1 and PC2). We ran a PCA with the metabolite classes as variables and explore the metabolomic differences amongst our four experimental groups in *Lasius niger* ants: **Y.F** (in green) = young foragers, **O.F** (in black) = old foragers, **Y.NW** (in blue) = young nest workers, **O.NW** (in red) = old nest-workers. The bar plots represent the classes of metabolites and their correlation with the first and second principal component (PC1 and PC2) from blue (negatively correlated) to red (positively correlated), as well as their contribution (bar length). All metabolites showed are correlated.

### 2.4. The emergence of anti-cancer mechanisms in the history of life

Smaller animals have fewer cells and therefore a lower probability of developing a tumor (as discussed in 66–68). These species therefore tend not to evolve mechanisms of cell replication blocking, found in bigger animals to prevent tumor development (66, 67). On the other hand, developing tumors over the course of life increases when the life span extends. Hence, long-lived social insects would face the situation of no replication-related protective mechanisms against cancer due to their size, combined with a higher probability of tumor development due to their longevity. However, no study describes a high prevalence of cancers among ants and eusocial insects in general. This raises the question of the mechanisms involved in the protection against cancer in those species. In our study, each group had at least one molecule potentially involved in anti-carcinogenic activity (e.g., [6]-shogaol, psilostachyin, thiophene; see **Table S8**) with a significantly higher relative abundance compared to the other groups. In other words, no clear variation due to age or social role could be identified for anti-carcinogenic molecules taken as a whole. The selection of molecular mechanisms of cancer resistance not based on replicative senescence is also found in naked mole rats, rodents that like ants are eusocial, small-sized, but long-lived (69).

Sociality, by increasing individual longevity, would therefore select for convergent anti-cancer mechanisms in distant taxa. However, only to a certain extent in ants since we found age-dependent abundances of proteins with pro-cancer activity, namely protein g12 and tetraspanin (70, 71). Old workers (O.F *vs* Y.F and O.NW *vs* Y.NW) had indeed larger amounts of these two proteins. Despite the presence of anti-cancer molecules, old ants might thus be more prone to cancer. This suggests that other age-related energetic trade-offs potentially made at the expense of anti-cancer mechanisms should also be taken into account. Still, it would also be possible that the observed amount of anti-cancer molecules co-varied in relation to other functions, which may have anti-aging effects, *e.g.*, immunity, involvement in the oxidative balance, antibiotic activity, or glycerophospholipid metabolism (see **Table S8**). The anti-cancer activity would be, in that case, only a side effect. Beyond the mechanistic questions, this would provide information on how sociality may impact cancer-related mechanisms, both at the intraspecific scale (workers vs queens, young vs old ants) and at the interspecific scale (between species of same or different level of sociality).

## 3. Conclusion

The purpose of this study was to disentangle the influence of social role and age on the physiology of workers from a eusocial species thanks to four experimental groups: young nest workers (Y.NW), young foragers (Y.F), old nest workers (O.NW) and old foragers (O.F). Specific functions (immunity, chemical communication, oxidative stress) and the clustering of samples showed in our study that both social role and age interact to define the individual phenotype.

Besides, our multi-omics approach highlighted evolutionary conserved mechanisms of aging (cancer, immunosenescence, poorer somatic maintenance in senescent groups, metabolic modifications). Thus, even if social insects, and more particularly ants, show distinctive features (notably the lack of a fertility/longevity trade-off), they share aging mechanisms with very distant taxa (mammals). This underlines potentially universal mechanisms, or at least shared by a majority of taxa and set-up very early in evolution of multicellular organisms (72). Furthermore, they offer us the opportunity to test how illness prevalence may be dependent on the synergy of ageing processes (73), under the influence of a strong degree of sociality. To describe more exhaustively the influence of sociality on individual physiology, further studies should include males and queens. In most ant species, males live only during the short breeding period, so it would be difficult to study the influence of age on their physiology. Conversely, because queens live up to ten times longer than workers (12), their regular monitoring over the course of their lifetime would take several years but should provide valuable information on the mechanisms slowing down aging processes. Beyond eusocial insects, uncovering mechanisms that promote a healthy lifespan could fuel progress in promoting human health.

## 4. Materials and Methods

The reader will find online (supplementary **Figure S1**) a graphical overview of the workflow applied in this study, from sample preparation of the four experimental groups, to the combined metabolomics and proteomics protocol and the curation of raw mass spectrometry data. Statistical analysis does not appear on this graphical overview but is detailed in the sections below.

### 4.1. Ant keeping and set-up of experimental groups

The black garden ant is a very common species in Western Europe where it predominantly inhabits urban habitats (74). For this species, the absence of dimorphism between workers (discussed in 75, 76) and the monogyny of the colonies reduce the potential sources of variation other than age and caste. In laboratory conditions, black garden ant workers have been shown to live 310 days on average and up to 1094 days, *ca.* 3 years (77). In our study, wild newly mated queen ants were collected in Lausanne, Switzerland (N 46.5234, E 6.5791). After being placed in individual glass tubes in the dark, 97 queens established a viable colony. Colonies were kept at a temperature of 21°C at night and 26°C during the day. To respect the biological rhythm of ants, we allowed them to enter diapause by gradually lowing the room temperature until 10°C for three months from December to March, before raising it to the usual values. The photoperiod varied throughout the year to mimic the natural photoperiod of the capture area. Relative humidity was 50-60%. Once a week, ant colonies were provided with a 0.3 M sugar water solution and mealworms *(Tenebrio molitor;* VenteInsecte, Courthézon, France).

After the first workers hatched, we let the colony develop for a month without any intervention, except feeding the ants. Due to the limited number of individuals available, young colonies at foundation sometimes have a less strict division of labour (78, 79). By waiting a month before the first collection, we allowed the colonies to grow sufficiently for the task specialization to take place. In addition, more developed colonies ensured that we could collect enough older individuals from the same colonies several months later. After a month, we ran the worker caste segregation protocol for the first time. The segregation of worker subcastes was based on their respective behaviors. Following a 48-hour period of starvation, colonies were proposed a 1 M sugar water. To maximize the forager recruitment, we waited five minutes after the first forager discovered the food source. Then, all the ants that came to the food source were considered as foragers, collected, and marked on the abdomen with an acrylic ink (Posca ©). We carried out this protocol three times to ensure we captured all foragers and with four-day intervals to allow the colony to rest. At the end of the procedure, not-marked ants were considered as nest workers. These young nest workers and foragers (0-1 month) were flash-frozen in liquid nitrogen and stored intact at −80°C until use. From that moment on, we followed the colonies carefully and removed the eggs in the last stages of maturation before they hatched. In this way, when we ran the worker caste segregation protocol for the second time, 11 months later, all the ants in the colonies were 11-12 months old. These two groups of old workers (nest workers and foragers) were also flash-frozen in liquid nitrogen and stored at −80°C until use. We ended up with four worker groups of both different behavior and age: young foragers (Y.F), old foragers (O.F), young nest worker (Y.NW) and old nest workers (O.NW). We sampled the experimental groups between August and September, several months before the ants enter diapause, which occurs from December to March in our laboratory conditions.

Before use for mass spectrometry analyses *(i.e.* proteomics and metabolomics), ink on foragers’ abdomens was removed with acetone. We constituted pools of 16 workers for each group and used 5 pools per group (80 workers per group). Pools were made of a balanced mixture from different colonies to remove this possible confounding effect (2-3 workers of each colony). Ants were ground under liquid nitrogen for 30s followed by 15s at 30 Hz with steel beads (Ø 2 mm, Mixer Mill MM400, Retsch, Eragny Sur Oise, France). Tubes containing the resulting powder and beads were then stored at −80°C until their use for mass spectrometry.

### 4.2. Proteomics analysis

Unless otherwise specified, all chemicals were purchased from Sigma Aldrich (St. Louis, MO, USA).

### 4.2.1. Sample Preparation

The frozen samples (powder) were suspended in 220 μL of lysis buffer (urea 8M, thiourea 2M, dithiothreitol [DTT] 1%, ammonium hydrogen carbonate [HNH_4_CO_3_] 0.1M, protease inhibitors 1.6 μM to 2 mM and sonicated on ice for 2 x 10 sec. at 135W, then samples were centrifuged (2000 x g, 2 min, RT) to eliminate possible cuticle remnants. Eight volumes of cold acetone (ThermoFisher Scientific, Rockford, IL, USA) were finally used for protein precipitation at −20°C overnight. Precipitated proteins were pelleted by centrifugation (13500 x g, 20 min, 4°C), washed once with cold acetone and then dissolved in Laemmli buffer (Tris 10 mM pH 7.5, EDTA 1 mM [Fluka, Buchs, Switzerland], β-mercaptoethanol 5%, SDS 5%, glycerol 10% [ThermoFisher Scientific]). Sonication and centrifugation were repeated as above to pellet and eliminate possibly remaining cell debris. Acetone fractions were all evaporated using a vacuum centrifuge (SpeedVac, Savant, Thermoscientific, Waltham, MA, USA) then kept at −20°C for further metabolomics analysis (see section *‘Metabolomics preparation’* below).

Total protein concentration was determined using the RC-DC Protein Assay kit (Bio-Rad, Hercules, CA, USA). At this stage, a reference sample comprising equal amounts of all protein extracts was made, to be analysed regularly during the whole experiment and allow QC-related measurements. Twenty micrograms of proteins from each sample were loaded onto SDS-PAGE gels (4% polyacrylamide for the stacking gel and 12% for the resolving gel) and electrophoresed for 20 minutes at 50 V then 20 minutes at 100 V. Proteins were thereafter fixed by a 15-minute incubation of gels in a solution composed of 50% ethanol and 3% phosphoric acid. Staining was performed using colloidal Coomassie Blue (30 min), and visualisation of proteins allowed five protein bands (2 mm each) to be excised from the gel. After destaining using acetonitrile/ammonium hydrogen carbonate 25 mM (75/25, v/v) and dehydration using pure acetonitrile, proteins were reduced and alkylated in-gel using 10 mM DTT in 25 mM ammonium hydrogen carbonate buffer (30 minutes at 60°C then 30 minutes at RT) and 55 mM iodoacetamide in 25 mM ammonium hydrogen carbonate buffer (20 min at RT in the dark), respectively. Gel slices were then washed using 25 mM ammonium hydrogen carbonate buffer (5 min, RT) and acetonitrile (5 min, RT) three times, and dehydration was finally performed using pure acetonitrile (2 x 5 min, RT). In-gel digestion of proteins was performed overnight at 37°C using trypsin (Promega Madison, WI, USA; 40 ng per band), and the resulting peptides were extracted twice (2 x 45 min) on an orbital shaker (450 rpm) using a solution composed of 60% acetonitrile and 0.5% formic acid in water. Another peptide extraction step was then performed (1 x 15 min) using 100% acetonitrile. At this stage, a set of reference peptides (iRT kit; Biognosys AG, Schlieren, Switzerland) was added to peptide extracts (6μL/sample after resuspension in 500mL of 20% acetonitrile/1% formic acid) for QC-related measurements. Organic solvent was thereafter evaporated using a vacuum centrifuge (SpeedVac) and the volume of peptide extracts was adjusted to 27 μL using a solution composed of 1% acetonitrile and 0.1% formic acid in water.

### 4.2.2. nanoLC-MS/MS analysis

NanoLC-MS/MS analysis was performed using a nanoUPLC-system (nanoAcquity; Waters, Milford, MA, USA) coupled to a quadrupole-Orbitrap hybrid mass spectrometer (Q-Exactive plus; Thermo Scientific, San Jose, CA, USA). The system was fully controlled by XCalibur software (v3.0.63; ThermoFisher Scientific). Samples (1 μl) were first concentrated/desalted onto a NanoEAse M/Z Symmetry precolumn (C18, 100 Å, 5 μm, 180 μm × 20 mm; Waters) using 99% of solvent A (0.1% formic acid in water) and 1% of solvent B (0.1% formic acid in acetonitrile) at a flow rate of 5 μl/min for 3 minutes. A solvent gradient from 1 to 6% of B in 0.5 minute then from 6 to 35% of B in 60 minutes was used for peptide elution, which was performed at a flow rate of 450 nL/min using a NanoEAse M/Z BEH column (C18, 130 Å, 1.7 μm, 75 μm x 250 mm; Waters) maintained at 60 °C. Samples were analysed randomly per block, each block being composed of one biological sample from each group. The reference sample was analysed six times throughout the experiment. In between each sample, washing of the column using 90% acetonitrile for 6 minutes and running of a solvent blank allowed limiting carry-over effects. Peak intensities and retention times of reference peptides were monitored in a daily fashion.

The Q-Exactive Plus was operated in positive ion mode with source temperature set to 250 °C and spray voltage to 1.8 kV. Full-scan MS spectra (300–1800 m/z) were acquired at a resolution of 70 000 at m/z 200. MS parameters were set as follows: maximum injection time of 50 ms, AGC target value of 3 × 10^6^ ions, lock-mass option enabled (polysiloxane, 445.12002 m/z), selection of up to 10 most intense precursor ions (doubly charged or more) per full scan for subsequent isolation using a 2 m/z window, fragmentation using higher energy collisional dissociation (HCD, normalised collision energy of 27), dynamic exclusion of already fragmented precursors set to 60 seconds. MS/MS spectra (300-2000 m/z) were acquired with a resolution of 17500 at m/z 200. MS/MS parameters were set as follows: maximum injection time of 100 ms, AGC target value of 1 × 10^5^ ions, peptide match selection option turned on.

### 4.2.3. Mass spectrometry data processing

Raw data were processed using MaxQuant v1.6.7.0 (80). Peak lists were created using default parameters. Using Andromeda search engine implemented in MaxQuant, peaklists were searched against a UniprotKb protein database *(Lasius niger,* TaxID 67767; 18217 entries) created in November 2019 with MSDA software suite (81). The database was complemented by Andromeda with the addition of the sequences of common contaminants like keratins and trypsin (247 entries) and of decoy (reverted) sequences for all *Lasius niger* proteins. Parameters were set as follows: precursor mass tolerance set to 20 ppm for the first search and to 4.5 ppm for the main search after recalibration, fragment ion mass tolerance set to 20 ppm, carbamidomethylation of cysteine residues considered as fixed modification, oxidation of methionines and acetylation of protein N-termini considered as variable modifications, peptide length of minimum 7 amino acids, maximum number of trypsin missed cleavages set to one, false discovery rate (FDR) set to 1% for both peptide spectrum matches and proteins. The proteins found with a single peptide or with a negative score were discarded from annotation data, as well as decoy hits and potential contaminants.

Protein quantification was performed using the MaxLFQ (label-free quantification) option implemented in MaxQuant. Parameters were set as follows: “minimal ratio count” of one, “match between runs” option enabled using a 0.7-minute time window after retention time alignment, consideration of both unmodified and modified (acetylation of protein N-termini and oxidation of methionines) peptides for quantitative determination, exclusion of shared peptides. All other MaxQuant parameters were set as default. Finally, criteria for retained proteins were as follows: at least two unique peptides quantified, no more than two missing values per group. Proteins absent in given groups (i.e. not detected at all) but satisfying above-mentioned criteria for the other groups were also retained. Among quantified proteins, 19 were annotated as “uncharacterized” (1.9% of all quantified proteins) for which we searched known homologous proteins in the Protostomia clade using BLAST searches (FASTA program v36; downloaded from http://fasta.bioch.virginia.edu/fasta_www2/fasta_down.shtml), and only the best hits were retained. To validate this procedure, we automatically extracted orthology annotations and sequence domains of *Lasius niger* uncharacterized proteins and of their homologues from the OrthoDB (82) and InterPRO (83) resources. The relevance of the match among *Lasius niger* uncharacterized proteins and their homologues was then checked manually. The mass spectrometry proteomics data have been deposited to the ProteomeXchange Consortium *via* the PRIDE (84) partner repository with the dataset identifier PXD026565. From QC-related measurements, we could see that the whole analysis system remained stable throughout the experiment. Indeed, a median coefficient of variation (CV) of 1.3% was calculated for retention times of iRT peptides over all injections, and a median CV of 28% was computed for all LFQ values obtained from the repeated analysis of the reference sample.

### 4.3. Metabolomics preparation

#### 4.3.1. Chemicals

Deionised water was filtered through a Direct-Q UV (Millipore) station, isopropanol and methanol were purchased from Fisher Chemicals (Optima ^®^ LC/MS grade). NaOH was obtained from Agilent Technologies, acetic acid, and formic acid from Sigma Aldrich.

#### 4.3.2. Sample preparation

Dried acetone fractions collected during sample preparation for proteomics (see above) were rehydrated with 500μl ethyl acetate and 300μl water. The samples were vortexed for 10 seconds, and the ethyl acetate phase was harvested after phase partitioning for each sample and stored until LC-MS/MS analysis. The water phase was also collected from each sample and diluted with 1 ml of water acidified with 1% formic acid. The acidified water phase was then desalted using Solid Phase Extraction (SPE) based on HLB matrix Oasis 96-Well plate (30 μm, 5mg; Waters) coupled to a vacuum pump. Each SPE well was conditioned with 1 ml of methanol, then with 1 ml of water. The samples were then applied on the SPE and washed with 1 ml of water acidified with 0.1% formic acid. The samples were then eluted with 700 μl of methanol. The elution fractions were mixed with the ethyl acetate fractions prior to further analysis and analyzed in liquid chromatography coupled to high resolution mass spectrometry (LC-HRMS) described in the next section.

#### 4.3.3. LC-HRMS analysis

Samples were analysed using liquid chromatography coupled to high resolution mass spectrometry on an UltiMate 3000 system (Thermo) coupled to an Impact II (Bruker) quadrupole time-of-flight (Q-TOF) spectrometer. Chromatographic separation was performed on an Acquity UPLC ^®^ BEH C18 column (2.1×100mm, 1.7 μm, Waters) equipped with and Acquity UPLC ^®^ BEH C18 pre-column (2.1×5mm, 1.7 μm, Waters) using a gradient of solvents A (Water, 0.1% formic acid) and B (MeOH, 0.1% formic acid). Chromatography was carried out at 35°C with a flux of 0.3mL.min^-1^, starting with 5% B for 2 minutes, reaching 100% B at 10 minutes, holding 100% for 3 minutes and coming back to the initial condition of 5% B in 2 minutes, for a total run time of 15 minutes. Samples were kept at 4°C, 10 μL were injected in full loop mode with a washing step after sample injection with 150 μL of wash solution (H_2_O/MeOH, 90/10, v/v). The spectrometer was operated in positive ion mode on a mass range of 20 to 1000 Da with a spectra rate of 2Hz in AutoMS/MS scan mode. The end plate offset was set to 500 V, capillary voltage at 2500 V, nebulizer at 2 Bar, dry gas at 8 L.min^-1^ and dry temperature at 200°C. The transfer time was set to 20-70μs and MS/MS collision energy at 80-120% with a timing of 50-50% for both parameters. The MS/MS cycle time was set to 3 seconds, absolute threshold to 816 cts and active exclusion was used with an exclusion threshold at 3 spectra, release after 1 min and precursor ion was reconsidered if the ratio current intensity/previous intensity was higher than 5. A calibration segment was included at the beginning of the runs allowing the injection of a calibration solution from 0.05 to 0.25min. The calibration solution used was a fresh mix of 50mL isopropanol/water (50/50, v/v), 500 μL NaOH 1M, 75 μL acetic acid and 25 μL formic acid. The spectrometer was calibrated in high precision calibration (HPC) mode with a standard deviation below 1 ppm before the injections, and recalibration of each raw data was performed after injection using the calibration segment.

#### 4.3.4. Metabolite annotation

Raw data were processed in MetaboScape 4.0 software (Bruker): molecular features were considered and grouped into buckets containing one or several adducts and isotopes from the detected ions with their retention time and MS/MS information when available. The parameters used for bucketing were a minimum intensity threshold of 10000, a minimum peak length of 4 spectra, a signal-to-noise ratio (S/N) of 3 and a correlation coefficient threshold set to 0.8. The [M+H]^+^, [M+Na]^+^ and [M+K]^+^ ions were authorized as possible primary ions, the [M+H-H2O] ion was authorized as common ion. Replicate samples were grouped and only the buckets found in 80% of the samples of at least one group were extracted from the raw data. The area of the peaks was used to compare the abundance of the features between the different groups. The obtained list of buckets was annotated using SmartFormula to generate raw formula based on the exact mass of the primary ions and the isotopic pattern. The maximum allowed variation on the mass (△m/z) was set to 3ppm, and the maximum mSigma value (assessing the good fitting of isotopic patterns) was set to 30. To put a name on the obtained formulae, metabolite lists were derived from Human Metabolite Database (HMDB, hmdb.ca), FooDB (foodb.ca), LipidMaps (lipidmaps.org) and SwissLipids (swisslipids.org). The parameters used for the annotation with the metabolite lists were the same as for SmartFormula annotation.

#### 4.4. Statistics and biological interpretation of mass spectrometry results

Unless otherwise specified, the analysis and graphical representations were made using R software, version 4.0 (85).

#### 4.4.1. Datasets

We only analyzed the proteins and metabolites (grouped under the term ‘analytes’), which we could consider present or absent in full confidence. Analytes were considered present in a group only when present in at least 3 out of 5 samples of this group. Conversely, an analyte was considered completely absent from a group only when none of the samples contained it. Consequently, analytes present in only 1 or 2 samples of a group were discarded from the statistical analysis. For analytes present in only 3 or 4 out of 5 samples of a given experimental group, we imputed the missing values using an iterative PCA (principal component analysis) algorithm (MissMDA package v.1.17, 86). Considering only the proteins and metabolites present or absent in a group according to our criteria, missing data represented 2.3% of the proteomics data and 0.9% of the metabolomic data. In the proteomics statistical analysis, we discarded one young forager sample and one young nest-worker sample identified as outliers during the statistical workflow. In all statistical analyses, proteins and metabolites were studied separately to better underline their respective roles in the physiology of ant workers.

#### 4.4.2. Heat maps and pairwise comparisons

First, we verified whether the different groups could be discriminated from each other using relative abundance of analytes and whether the discrimination criterion was age or behavior. For this, we used the ‘rlog’ function of the DESeq2 package (v.1.28, 87) to transform the proteomics and metabolomics data to the log2 scale in a way which minimizes differences among samples with small counts, and normalizes with respect to the size of the dataset (88). Based on this normalized data, we built heat maps with the ComplexHeatmap package (v.2.42, 89). Heat maps represent the most differentially expressed analytes and perform a hierarchical clustering, which reflects the proteomic or metabolomic similarities among samples. Then, we sought to identify exactly which proteins and which metabolites were expressed depending on the age or behavior. Comparing O. NW vs. O.F or Y.NW vs. Y.F provides information on the influence of the behavior since the age among the groups compared is identical. Conversely, keeping the behavior factor constant, by comparing Y.NW vs O.NW or Y.F vs O.F, makes it possible to assess the influence of age. We identified the analytes that differed most strongly between the groups compared two-by-two by calculating fold-changes (FC) of each analyte with the DESeq2 package. In this analysis, we retained only the analytes with a false discovery rate (FDR) lower than 0.05 and a FC higher than 2 (up-regulated) or lower than −2 (down-regulated).

#### 4.4.3. Classification and functional annotation of analytes

In metabolomics, a molecule is stated *‘identified’* only when compared with a reference standard. However, we did not use a standard for the hundreds of metabolites found in this study, but we used online databases (see sections above) to attribute names to the analytes found. This is called annotation: in this study, the presented annotations are at the level 3 of Schymanski’s classification (90). Conversely, peptides do not require comparison to a standard to be formally identified. However, for consistency throughout the article, we used the term *“annotated”* to referred to *annotated* metabolites and *identified* proteins. Consequently, when an analyte (protein or metabolite) is simply referred to as *‘annotated’,* it means named and not functionally annotated.

To address the function of detected molecules, we ran functional enrichment analyses. Such methods evaluate the significance of a set of functionally related molecules. Regarding the proteome, functional annotation enrichment analysis was performed using the desktop version of DAVID (Ease v2.1) and versions of Gene Ontology (GO) and KEGG databases downloaded in December 2021. However, no such annotation was available for ant proteins, and no significant enrichment has been found when using homologous proteins in *Drosophila melanogaster* or *Homo sapiens.* Regarding the metabolome, we used the online platform MetaboAnalyst (v. 5.0, www.metaboanalyst.ca, 91) and performed a metabolite set enrichment analysis (aka. MSEA). However, among 196 annotated metabolites kept for the characterization of the four experimental worker groups, only 53 (27.04%) were recognized by the platform. This issue presumably came from the fact that the databases used by MetaboAnalyst are mainly derived from human studies. The analysis did not sufficiently cover our dataset to fully depict its diversity. The results of this analysis will therefore not be discussed but are available as supplementary material (**ESM2**). As automated functional enrichment was not compatible with our dataset, we investigated the biological meaning of the proteomics and metabolomics profiles by automatically retrieving metabolic maps from the KEGG database (genome.jp/kegg) when available. If not, we “manually” looked in the literature for functions fulfilled by the molecules in concern. References are provided in Tables S7 and S8.

Metabolites may encompass very distinct molecules, *e.g.,* lipids, free amino acids, free nucleic acids, carbohydrates. We automatically classified metabolites thanks to the ChemRICH online tool (chemrich.fiehnlab.ucdavis.edu; 92), which covered 79.03% of our metabolite dataset. We used these classes of metabolites in a principal component analysis (PCA, **Figure 3**) to see whether some groups of metabolites characterized rather the behavior than age, and conversely (FactoMineR package v.2.3, 93). Using the PCA coordinates, we calculated the intra-group repeatability (package “rptR” v.09.22, 94) to assess their homogeneity. The metabolite’s class was also added to heat maps to provide additional information about the most segregating molecules.

## Supporting information

Data sets

## 5. Abbreviations

FC: fold change
FDR: false discovery rate
PC (A): principal component (analysis)
Y/O.NW: young/old nest-workers
Y/O.F: young/old foragers

## 6. Acknowledgments

We thank Nathalie Stroeymeyt for providing the newly-mated queens, Hélène Gachot-Neveu, Aurélie Kranitsky and David Bock for their precious work in the animal husbandry.

The study was supported by the Centre National de la Recherche Scientifique (CNRS) and the French Proteomic Infrastructure (ProFi; ANR-10-INSB-08-03). M. Quque and C. Brun PhDs were funded by the University of Strasbourg and the Ministère de l’Enseignement Supérieur, de la Recherche et de l’Innovation (French Ministry of higher Education, Research and Innovation).

## References

1. C. Sueur, M. Quque, A. Naud, A. Bergouignan, F. Criscuolo, Social capital: an independent dimension of healthy ageing. Peer Community Journal 1 (2021).

2. R. Covas, M. A. du Plessis, C. Doutrelant, Helpers in colonial cooperatively breeding sociable weavers *Philetairus socius* contribute to buffer the effects of adverse breeding conditions. Behav Ecol Sociobiol 63, 103–112 (2008).

3. D. Aydinonat, et al., Social isolation shortens telomeres in African grey parrots (Psittacus erithacus erithacus). PLoS ONE 9, e93839 (2014).

4. H. Ruan, C.-F. Wu, Social interaction-mediated lifespan extension of Drosophila Cu/Zn superoxide dismutase mutants. PNAS 105, 7506–7510 (2008).

5. E. H. Dawson, et al., Social environment mediates cancer progression in Drosophila. Nature Communications 9 (2018).

6. B. Wild, et al., Social networks predict the life and death of honey bees. bioRxiv, 2020.05.06.076943 (2020).

7. N. Snyder-Mackler, et al., Social determinants of health and survival in humans and other animals. Science 368, eaax9553 (2020).

8. I. Kawachi, S. V. Subramanian, D. Kim, “Social Capital and Health” in Social Capital and Health, I. Kawachi, S. V. Subramanian, D. Kim, Eds. (Springer, 2008), pp. 1–26.

9. N. Grant, M. Hamer, A. Steptoe, Social isolation and stress-related cardiovascular, lipid, and cortisol responses. Ann Behav Med 37, 29–37 (2009).

10. L. Keller, S. Jemielity, Social insects as a model to study the molecular basis of ageing. Experimental gerontology 41, 553–556 (2006).

11. J. D. Parker, What are social insects telling us about aging? Myrmecological News 13, 103–110 (2010).

12. L. Keller, M. Genoud, Extraordinary lifespans in ants: a test of evolutionary theories of ageing. Nature 389, 958–960 (1997).

13. J. D. Parker, K. M. Parker, B. H. Sohal, R. S. Sohal, L. Keller, Decreased expression of Cu-Zn superoxide dismutase 1 in ants with extreme lifespan. Proc Natl Acad Sci U S A 101, 101, 3486, 3486–3489 (2004).

14. M. Corona, K. A. Hughes, D. B. Weaver, G. E. Robinson, Gene expression patterns associated with queen honey bee longevity. Mechanisms of Ageing and Development 126, 1230–1238 (2005).

15. M. Corona, G. E. Robinson, Genes of the antioxidant system of the honey bee: annotation and phylogeny. Insect Molecular Biology 15, 687–701 (2006).

16. E. R. Lucas, L. Keller, Ageing and somatic maintenance in social insects. Current Opinion in Insect Science 5, 31–36 (2014).

17. B. J. Heidinger, et al., Telomere length in early life predicts lifespan. PNAS 109, 1743-1748 (2012).

18. J. R. Eastwood, et al., Early-life telomere length predicts lifespan and lifetime reproductive success in a wild bird. Molecular Ecology 28, 1127–1137 (2018).

19. K. Whittemore, E. Vera, E. Martínez-Nevado, C. Sanpera, M. A. Blasco, Telomere shortening rate predicts species life span. PNAS, 201902452 (2019).

20. S. Jemielity, et al., Short telomeres in short-lived males: what are the molecular and evolutionary causes? Aging Cell 6, 225–233 (2007).

21. I. Iovinella, et al., Antennal protein profile in honeybees: caste and task matter more than age. Front. Physiol. 9 (2018).

22. C. W. Whitfield, A. M. Cziko, G. E. Robinson, Gene expression profiles in the brain predict behavior in individual honey bees. Science 302, 296–299 (2003).

23. A. Dussutour, L.-A. Poissonnier, J. Buhl, S. J. Simpson, Resistance to nutritional stress in ants: when being fat is advantageous. Journal of Experimental Biology 219, 824–833 (2016).

24. P. Kohlmeier, A. R. Alleman, R. Libbrecht, S. Foitzik, B. Feldmeyer, Gene expression is more strongly associated with behavioural specialisation than with age or fertility in ant workers. Molecular Ecology (2018) https:/doi.org/10.1111/mec.14971 (December 10, 2018).

25. E. R. Lucas, J. Romiguier, L. Keller, Gene expression is more strongly influenced by age than caste in the ant Lasius niger. Mol Ecol 26, 5058–5073 (2017).

26. C. Alaux, et al., Regulation of brain gene expression in honey bees by brood pheromone. Genes, Brain and Behavior 8, 309–319 (2009).

27. Y. B. Jarrar, S.-J. Lee, Molecular functionality of cytochrome P450 4 (CYP4) genetic polymorphisms and their clinical implications. International Journal of Molecular Sciences 20, 4274 (2019).

28. M. J. Snyder, Cytochrome P450 enzymes belonging to the CYP4 family from marine invertebrates. Biochemical and Biophysical Research Communications 249, 187–190 (1998).

29. R. Feyereisen, Insect P450 Enzymes. Annual Review of Entomology 44, 507–533 (1999).

30. E.-J. Won, et al., Expression of three novel cytochrome P450 (CYP) and antioxidative genes from the polychaete, *Perinereis nuntia* exposed to water accommodated fraction (WAF) of Iranian crude oil and Benzo[α]pyrene. Marine Environmental Research 90, 75-84 (2013).

31. D. L. Geiser, J. J. Winzerling, Insect transferrins: Multifunctional proteins. Biochimica et Biophysica Acta (BBA) - General Subjects 1820, 437–451 (2012).

32. T. Obuchi, et al., Armillaric acid, a new antibiotic produced by *Armillaria mellea*. Planta medica 56, 198–201 (1990).

33. J. Šmidrkal, T. Karlová, V. Filip, M. Zárubová, I. Hrádková, Antimicrobial properties of 11-cyclohexylundecanoic acid. Czech Journal of Food Sciences 27 (2009), 463–469 (2009).

34. G. W. Felton, C. B. Summers, Antioxidant systems in insects. Archives of Insect Biochemistry and Physiology 29, 187–197 (1995).

35. K. S. Lee, et al., Transferrin inhibits stress-induced apoptosis in a beetle. Free Radical Biology and Medicine 41, 1151–1161 (2006).

36. H. Ikeda, Y. Togashi, Aging, cancer, and antitumor immunity. Int J Clin Oncol 27, 316–322 (2022).

37. L. S. Haddad, L. Kelbert, A. J. Hulbert, Extended longevity of queen honey bees compared to workers is associated with peroxidation-resistant membranes. Experimental Gerontology 42, 601–609 (2007).

38. R. Pamplona, G. Barja, M. Portero-Otín, Membrane fatty acid unsaturation, protection against oxidative stress, and maximum life span. Annals of the New York Academy of Sciences 959, 475–490 (2002).

39. A. J. Hulbert, R. Pamplona, R. Buffenstein, W. A. Buttemer, Life and death: metabolic rate, membrane composition, and life span of animals. Physiological Reviews 87, 1175–1213 (2007).

40. M. Chapuisat, L. Keller, Division of labour influences the rate of ageing in weaver ant workers. Proceedings of the Royal Society of London B: Biological Sciences 269, 909–913 (2002).

41. P. Kohlmeier, et al., Intrinsic worker mortality depends on behavioral caste and the queens’ presence in a social insect. The Science of Nature 104 (2017).

42. M. J. Toth, A. Tchernof, Lipid metabolism in the elderly. Eur J Clin Nutr 54, S121–S125 (2000).

43. L. I. Gilbert, “Lipid Metabolism and Function in Insects” in Advances in Insect Physiology, (Elsevier, 1967), pp. 69–211.

44. D. W. Stanley-Samuelson, R. A. Jurenka, C. Cripps, G. J. Blomquist, M. de Renobales, Fatty acids in insects: Composition, metabolism, and biological significance. Arch. Insect Biochem. Physiol. 9, 1–33 (1988).

45. K. H. Lockey, Lipids of the insect cuticle: origin, composition and function. Comparative Biochemistry and Physiology Part B: Comparative Biochemistry 89, 595–645 (1988).

46. B. J. Sinclair, K. E. Marshall, The many roles of fats in overwintering insects. Journal of Experimental Biology 221, jeb161836 (2018).

47. Q. Sun, J. Li, F. Gao, New insights into insulin: The anti-inflammatory effect and its clinical relevance. World J Diabetes 5, 89–96 (2014).

48. M. A. Cooper, et al., Reduced mitochondrial reactive oxygen species production in peripheral nerves of mice fed a ketogenic diet. Experimental Physiology 103, 1206–1212 (2018).

49. C. Xu, et al., Feeding restriction alleviates high carbohydrate diet-induced oxidative stress and inflammation of *Megalobrama amblycephala* by activating the AMPK-SIRT1 pathway. Fish & Shellfish Immunology 92, 637–648 (2019).

50. M. B. Blackburn, M. J. Loeb, E. Clark, H. Jaffe, Stimulation of midgut stem cell proliferation by *Manduca sexta* α-arylphorin. Archives of Insect Biochemistry and Physiology 55, 26–32 (2004).

51. R. S. Hakim, et al., Growth and mitogenic effects of arylphorin in vivo and in vitro. Archives of Insect Biochemistry and Physiology 64, 63–73 (2007).

52. A. Castagnola, et al., Alpha-arylphorin is a mitogen in the Heliothis virescens midgut cell secretome upon Cry1Ac intoxication. PeerJ 5, e3886 (2017).

53. M. Quque, et al., Division of labour in the black garden ant *(Lasius niger)* leads to three distinct proteomes. Journal of Insect Physiology 117, 103907 (2019).

54. M. Quque, et al., Eusociality is linked to caste-specific differences in metabolism, immune system, and somatic maintenance-related processes in an ant species. Cell. Mol. Life Sci. 79, 29 (2021).

55. C. Buehlmann, P. Graham, B. S. Hansson, M. Knaden, Desert ants locate food by combining high sensitivity to food odors with extensive crosswind runs. Current Biology 24, 960–964 (2014).

56. H.-L. Qiu, D.-F. Cheng, A chemosensory protein gene Si-CSP1 associated with necrophoric behavior in red imported fire ants (hymenoptera: formicidae). J Econ Entomol 110, 1284-1290 (2017).

57. Q. Sun, K. F. Haynes, X. Zhou, Managing the risks and rewards of death in eusocial insects. Philosophical Transactions of the Royal Society B: Biological Sciences 373, 20170258 (2018).

58. D. M. Gordon, Dependence of necrophoric response to oleic acid on social context in the ant, Pogonomyrmex badius. J Chem Ecol 9, 105–111 (1983).

59. Q. Sun, X. Zhou, Corpse management in social insects. Int J Biol Sci 9, 313–321 (2013).

60. L. Diez, L. Moquet, C. Detrain, Post-mortem changes in chemical profile and their influence on corpse removal in ants. J Chem Ecol 39, 1424–1432 (2013).

61. Q. Sun, K. F. Haynes, X. Zhou, Dynamic changes in death cues modulate risks and rewards of corpse management in a social insect. Functional Ecology 31, 697–706 (2017).

62. S. Cremer, S. A. O. Armitage, P. Schmid-Hempel, Social immunity. Current Biology 17, R693–R702 (2007).

63. N. Stroeymeyt, et al., Social network plasticity decreases disease transmission in a eusocial insect. Science 362, 941–945 (2018).

64. J. Heinze, B. Walter, Moribund ants leave their nests to die in social isolation. Current Biology 20, 249–252 (2010).

65. N. Bos, T. Lefèvre, A. B. Jensen, P. D’Ettorre, Sick ants become unsociable. Journal of Evolutionary Biology 25, 342–351 (2012).

66. A. Seluanov, et al., Distinct tumor suppressor mechanisms evolve in rodent species that differ in size and lifespan. Aging Cell 7, 813–823 (2008).

67. V. Gorbunova, A. Seluanov, Coevolution of telomerase activity and body mass in mammals: From mice to beavers. Mechanisms of Ageing and Development 130, 3–9 (2009).

68. X. Tian, et al., Evolution of telomere maintenance and tumour suppressor mechanisms across mammals. Philosophical Transactions of the Royal Society B: Biological Sciences 373, 20160443 (2018).

69. A. Seluanov, V. N. Gladyshev, J. Vijg, V. Gorbunova, Mechanisms of cancer resistance in long-lived mammals. Nature Reviews Cancer 18, 433–441 (2018).

70. J. Juneja, P. J. Casey, Role of G12 proteins in oncogenesis and metastasis. British Journal of Pharmacology 158, 32–40 (2009).

71. H.-X. Wang, Q. Li, C. Sharma, K. Knoblich, M. E. Hemler, Tetraspanin protein contributions to cancer. Biochemical Society Transactions 39, 547–552 (2011).

72. C. A. Aktipis, et al., Cancer across the tree of life: cooperation and cheating in multicellularity. Philosophical Transactions of the Royal Society B: Biological Sciences 370, 20140219 (2015).

73. J.-F. Lemaître, et al., Eco-evolutionary perspectives of the dynamic relationships linking senescence and cancer. Functional Ecology 34, 141–152 (2020).

74. E. A. Konorov, et al., Genomic exaptation enables *Lasius niger* adaptation to urban environments. BMC Evolutionary Biology 17, 39 (2017).

75. I. M. Grześ, M. Okrutniak, J. Grzegorzek, The size-dependent division of labour in monomorphic ant Lasius niger. European Journal of Soil Biology 77, 1–3 (2016).

76. M. Okrutniak, B. Rom, F. Turza, I. M. Grześ, Body size differences between foraging and intranidal workers of the monomorphic ant Lasius niger. Insects 11, 433 (2020).

77. B. H. Kramer, R. Schaible, A. Scheuerlein, Worker lifespan is an adaptive trait during colony establishment in the long-lived ant Lasius niger. Exp. Gerontol. 85, 18–23 (2016).

78. C. R. Ferreira Brandao, Sequential ethograms along colony development of *Odontomachus affinis* Guérin (Hymenoptera, Formicidae, Ponerinae). Ins. Soc 30, 193-203 (1983).

79. C. T. Holbrook, P. M. Barden, J. H. Fewell, Division of labor increases with colony size in the harvester ant Pogonomyrmex californicus. Behav Ecol 22, 960–966 (2011).

80. J. Cox, et al., Accurate Proteome-wide Label-free Quantification by Delayed Normalization and Maximal Peptide Ratio Extraction, Termed MaxLFQ. Mol Cell Proteomics 13, 2513–2526 (2014).

81. C. Carapito, et al., MSDA, a proteomics software suite for in-depth Mass Spectrometry Data Analysis using grid computing. Proteomics 14, 1014–1019 (2014).

82. E. V. Kriventseva, et al., OrthoDB v10: sampling the diversity of animal, plant, fungal, protist, bacterial and viral genomes for evolutionary and functional annotations of orthologs. Nucleic Acids Res. 47, D807–D811 (2019).

83. A. L. Mitchell, et al., InterPro in 2019: improving coverage, classification and access to protein sequence annotations. Nucleic Acids Res. 47, D351–D360 (2019).

84. J. A. Vizcaíno, et al., 2016 update of the PRIDE database and its related tools. Nucleic acids research 44, D447–D456 (2016).

85. R Core Team, R: A language and environment for statistical computing. (R Foundation for Statistical Computing, 2019).

86. J. Josse, F. Husson, missMDA: a package for handling missing values in multivariate data analysis. Journal of Statistical Software 70, 1–31 (2016).

87. M. Love, Assessment of DESeq2 performance through simulation. www.huber.embl.de/DESeq2paper (2014) (April 4, 2019).

88. M. I. Love, W. Huber, S. Anders, Moderated estimation of fold change and dispersion for RNA-seq data with DESeq2. Genome Biology 15, 550 (2014).

89. Z. Gu, R. Eils, M. Schlesner, Complex heatmaps reveal patterns and correlations in multidimensional genomic data. Bioinformatics 32, 2847–2849 (2016).

90. E. L. Schymanski, et al., Identifying small molecules via high resolution mass spectrometry: communicating confidence. Environ. Sci. Technol. 48, 2097–2098 (2014).

91. Z. Pang, et al., MetaboAnalyst 5.0: narrowing the gap between raw spectra and functional insights. Nucleic Acids Research 49, W388–W396 (2021).

92. D. K. Barupal, O. Fiehn, Chemical similarity enrichment analysis (ChemRICH) as alternative to biochemical pathway mapping for metabolomic datasets. Scientific Reports 7, 14567 (2017).

93. S. Lê, J. Josse, F. Husson, FactoMineR: an R package for multivariate analysis. Journal of Statistical Software 025, 1–18 (2008).

94. M. A. Stoffel, S. Nakagawa, H. Schielzeth, rptR: repeatability estimation and variance decomposition by generalized linear mixed-effects models. Methods Ecol Evol 8, 1639-1644 (2017).

