## Supplementary Materials for "Both social environment and chronological age shape the physiology of ant workers"

Electronic supplementary Material

Sociality shapes aging and physiology: the multi-omics case of the ant workers.

**ESM1 – Workflow of the joint proteomics-metabolomics analysis**

**[below] Figure S1. Workflow from sample preparation to data analysis.** **A)** After collecting freshly mated queens of black garden ants (*Lasius niger*) in the field, we let new colonies to settle during one month, then we collected young foragers (*Y.F*) and young nest-workers (*Y.NW*). From this date, larvae were removed and 11 months later, we collected workers again: old foragers (*O.F*) and old nest-workers (*O.NW*). Samples were stored intact at -80°C and processed at the same time in the mass spectrometry analyses. **B)** The proteomics (at left in green) and metabolomics (at right in red) protocols were run in parallel as shown in this figure and detailed in the “Material and Methods” section. **C)** We detected 1719 proteins and 712 metabolites among the four experimental groups. An analyte (metabolite or protein) was considered as ‘*present in all castes*’ when found in at least 3 samples per caste and as ‘*absent in at least one caste*’ when not found in any sample of one or more caste. All other analytes were *not selected* for further analysis. An *annotated* analyte means that a name was automatically attributed by querying databases. Further details in main text.

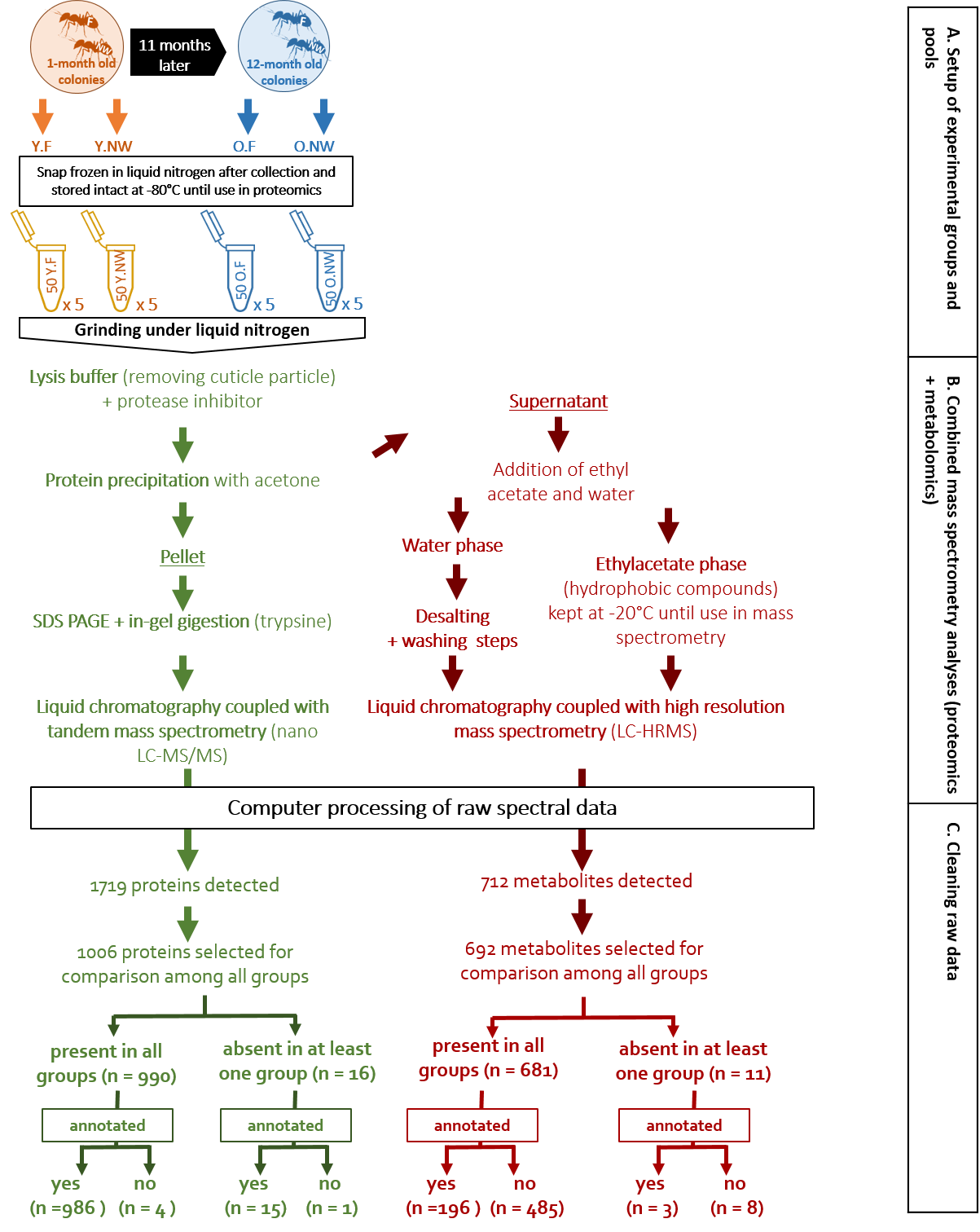

### ESM2 – Functional enrichment Analysis

To understand the biological meaning of proteomics and metabolomics profiles, we ran functional enrichment analysis. This method compares the relative expression of metabolic pathways, according to amount of molecules involved in them, among experimental groups. Thus, we aimed to retrieve a functional profile of our experimental groups.

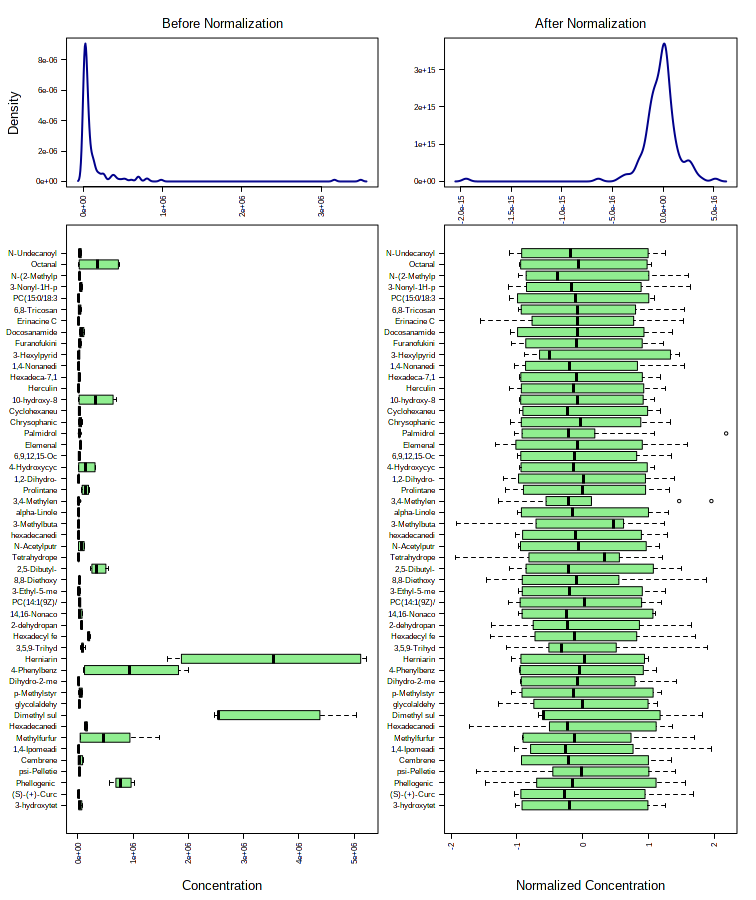

#### Metabolomics

**Figure S2 Normalization of our metabolomics dataset.** Figures provided by MetaboAnalyst after normalization through the ‘auto-scaling’ option (mean-centered and divided by the standard deviation of each variable).

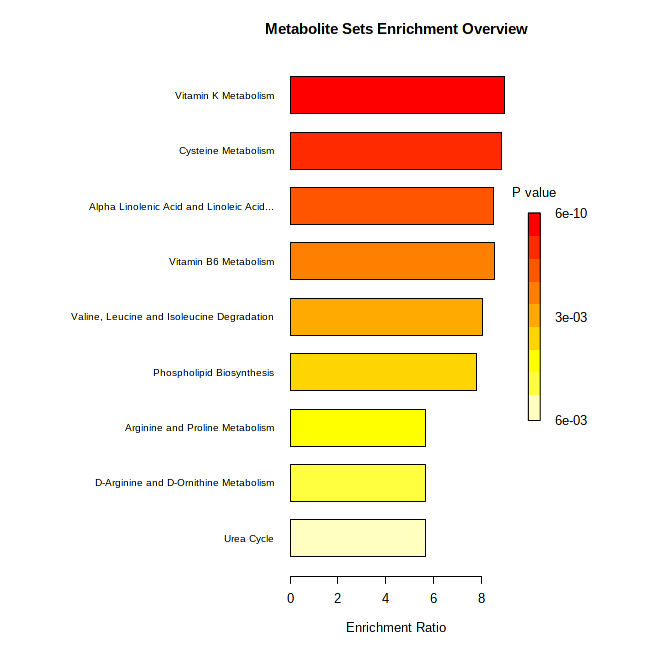
We used the online platform MetaboAnalyst (v. 5.0, www.metaboanalyst.ca; Pang et al. 2021) and ran an automated metabolite set enrichment analysis (aka. MSEA). Metabolites are usually assessed individually for their significance under the study conditions. On the contrary, MSEA directly evaluates the significance of a set of functionally related metabolites. During the quantitative enrichment analysis, the data were normalized with the ‘auto-scaling’ option (see **Figure S2** above). We queried both Small Molecule Pathway Database (SMPDB) and Kyoto Encyclopedia of Genes and Genomes (KEGG), since they provide complementary pathways. We considered all molecules in these databases that had at least two entries. However, the databases used by MetaboAnalyst are mainly derived from human studies. This led to the fact that among 196 annotated metabolites kept for the characterization of the four experimental worker groups, only 53 (27.04%) were recognized by the platform. The databases used by the platform do not sufficiently cover our dataset to fully depict its diversity. Outcomes of the analysis can be found below.

**Figure S3 Metabolite Set Enrichment using SMPDB**

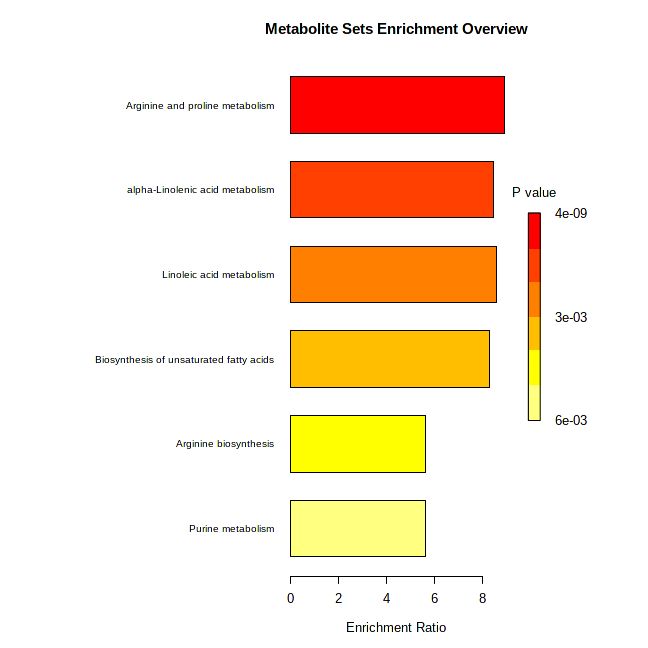

**Figure S4 Metabolite Set Enrichment using KEGG**

We can notice overlaps between both databases. The quantitative enrichment analyses mostly highlighted pathways linked to amino acids (arginine, valine, leucine), linoleic acids, and vitamins. As we can notice from the comparison with the methodology used in main text, using this approach would have led to miss many biological functions (e.g., oxidative status, digestive function, cancer-related molecules). As discussed above, this is probably linked to the fact that those database are intended for studies in humans.
